# Autistic traits modulate neural responses to social signals during natural vision

**DOI:** 10.1101/2025.10.16.682799

**Authors:** Qingying Ye, Jinglu Chen, Severi Santavirta, Vesa Putkinen, Juha Salmi, Lauri Nummenmaa

**Author notes:** Corresponding author: Qingying Ye Postal address: Turku PET Centre, 4-8 Kiinamyllynkatu, 20520 Turku Contact.

## Abstract

Impairments in social perception, a hallmark of autism spectrum disorder (ASD), are also evident at subclinical levels in the general population. However, it remains unclear how such variation in autistic traits modulate neural processing of different types of social information. Here, we investigated whether autistic traits in neurotypical individuals are associated with neural responses to a broad array of social perceptual features during viewing naturalistic stimuli using functional magnetic resonance imaging (fMRI). We also tested the generalizability of these effects across two experiments. Ninety-seven participants completed the Autism Spectrum Quotient (AQ) and watched a set of 96 movie clips and a full movie during an fMRI scan. Intensity of 126 social features in the movie stimuli was continuously annotated by independent observers, and 44 most reliably rated features were used to model neural responses. We examined how consistently the responses to each social feature were dependent on the participant’s AQ scores. Replicable AQ-dependent neural responses to social features were found in both datasets. The temporal cortex and especially the superior temporal gyrus (STG), served as a central “hub” where autistic traits consistently modulated responses to social features across datasets. Different AQ subscales also revealed distinct association patterns in other brain regions. These findings indicate that autism-related traits broadly influence neural processing of naturalistic social signals, providing insight into how characteristics of autistic symptoms relate to socioemotional processing.

## 1 Introduction

Autism spectrum disorder (ASD) is a heterogeneous neurodevelopmental condition characterized by impairments in social interaction and communication, along with repetitive behaviors and restricted interests (American Psychiatric Association, 2013). Subclinical variation in similar behavioral characteristics is observed in the general population, and such autism-related traits are conceptualized as phenotype markers (Constantino et al., 2003). Individuals with higher levels of autistic traits show impaired performance on social cognition tasks such as perception of social cues (Alkhaldi et al., 2024), emotional recognition (Corluka and Laycock, 2024), and mentalizing (Le et al., 2024), resembling the patterns observed in ASD. Therefore, studying these subclinical traits with neuroimaging methods in the general population could elucidate the brain basis of autism-related impairments while minimizing the confounding effects of disease course, chronicity, medication, and co-morbid conditions often present in clinical studies.

Aberrant social cognition, especially deficits in social perception, is one of the key factors behind social difficulties in ASD. Social perception refers to the early stage of sensory information processing in which individuals detect and interpret cues critical for understanding others’ intentions and dispositions (Allison et al., 2000). Various aspects of social perception in ASD have been investigated at behavioral (Chita-Tegmark, 2016; Riddiford et al., 2022) and neural (Deen et al., 2015; Floris et al., 2025; Limanowski and Blankenburg, 2016; Ropar et al., 2018) levels. For example, eye-tracking meta-analyses show that individuals with ASD exhibit reduced eye gaze toward social stimuli, particularly the eye region of the human face, compared to neurotypical controls (Chita-Tegmark, 2016; Riddiford et al., 2022). Neuroimaging studies have revealed structural and functional impairments in ASD across brain regions involved in perception of eye contact (superior temporal sulcus, STS; Pelphrey et al., 2005), faces (fusiform gyrus; Floris et al., 2025), bodies (temporo-parietal junction, TPJ; Limanowski and Blankenburg, 2016; Ropar et al., 2018), and other higher-order social-cognitive processes (e.g., superior temporal gyrus, STG; medial prefrontal cortex, mPFC; Amodio and Frith, 2006; Redcay, 2008).

The above evidence suggests that ASD is associated with neural alterations across a broad range of social perceptual domains. Similar neural alterations are also observed in the general populations with autistic traits during perceiving eye gaze (Nummenmaa et al., 2012a), social voice (Skjegstad et al., 2022), and social touch (Voos et al., 2013). However, research on social perception has typically used simplified study designs (e.g., static images) and studied isolated social features one at a time, although it is unlikely that social features are processed independently of other social information by distinct neural regions or systems (Huth et al., 2012; van den Heuvel and Sporns, 2011). Therefore, evidence from studies using simplified stimuli may not generalize to real-world social perception.

Movie-watching combined with functional magnetic resonance imaging (fMRI) enhances ecological validity, yielding results that are more generalizable to real-world settings and demonstrating significant advantages in research on social perception. Movie stimuli provide continuous, naturalistic input with rich social and emotional contents, eliciting cognitive and affective processes that closely approximate real-world experience (Hasson et al., 2004; Nummenmaa et al., 2012b; Salmi et al., 2013). fMRI studies using movie viewing paradigms have revealed a distributed but functionally organized neural network for social perception among the general population, highlighting the STS as a general hub for social perception (Lahnakoski et al., 2012). Furthermore, this network follows a posterior-to-anterior gradient: temporal, occipital and parietal regions are broadly engaged in the perception of diverse social features, whereas frontal and subcortical regions are recruited more selectively (Santavirta et al., 2023). Additionally, the neural basis of separate social features (e.g., laughing, crying, and biological motion) in natural scenes has also been investigated in the general population (Hudson et al., 2023; Nummenmaa et al., 2023). Regarding autism-like traits, one study reported that individuals with ASD have reduced activation compared to controls in areas related to mentalizing and perspective taking in the right TPJ and right posterior STS when watching socially awkward episodes (Pantelis et al., 2015). Moreover, another study found that autistic traits were associated with decreased neural activity in middle cingulate gyrus and precuneus during the perception of naturalistic biological motion (Hudson et al., 2023).

### The Current Study

Despite the growing literature on the neural basis of social perception in general population, limited research has focused on studying how autistic traits modulate neural responses for social perception under naturalistic conditions. Since most previous studies have mapped how autistic traits relate to the perception of isolated social features although, in reality, multiple social features are perceived simultaneously, a more comprehensive understanding of how these traits affect neural responses to diverse aspects of social perception is needed. Here, we tested whether autistic traits in neurotypical populations are associated with neural responses when viewing naturalistic movies in two experiments with independent stimulus sets. Participants (N=97) completed the Autism Spectrum Quotient (AQ), and their haemodynamic brain activity was measured with fMRI when viewing a set of 96 movie clips (**Movie clips** dataset) and a full movie (**Full movie** dataset, see **Figure 1**). The presence of 126 social features was dynamically rated in a separate experiment by independent observers to get a comprehensive description of the social contents. The 44 most consistently evaluated social features were then selected for analyses, to enable reliable modelling of the neural responses in the independent fMRI sample (Santavirta et al., 2023). We then tested how consistently the responses to each social feature were dependent on the participant’s AQ scores, both the total score and subscales. The present results indicated both general and dimension-specific associations between AQ scores and neural responses to social features in two datasets, suggesting that autism-like traits consistently modulate the neural processing of naturalistic audiovisual social information in the general population.

**Figure 1.**
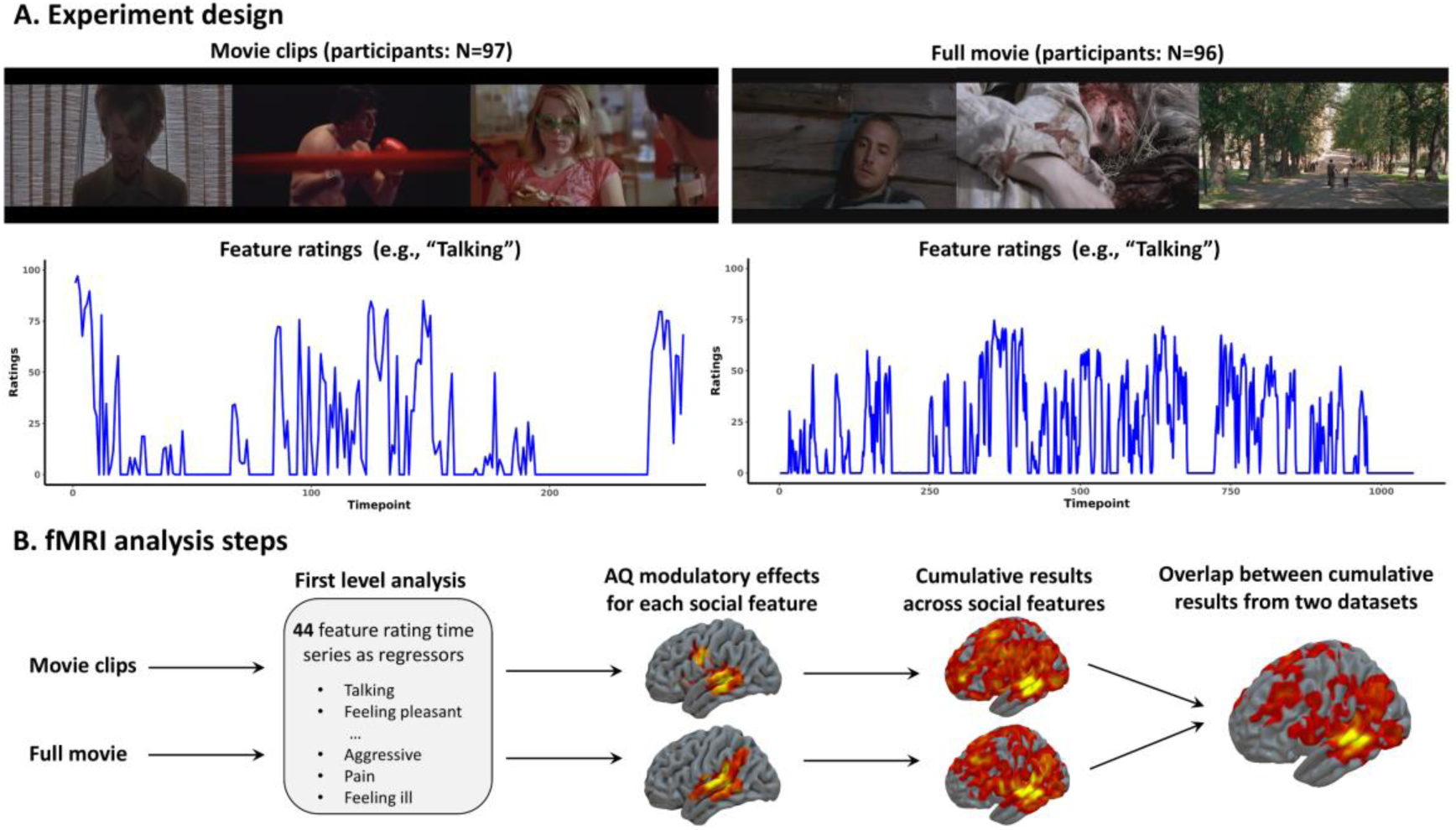
Experimental design and analytical pipeline of the study. (A) Participants (N=97) watched 96 unrelated video clips of social scenes and a full movie in two separate fMRI sessions. (B) Social contents of the stimuli were dynamically annotated and 44 most consistently annotated social feature ratings were used to model the heamodynamic responses of neurotypical participants. Next, their AQ scores were used to model the interindividual differences in the response patterns for each social feature. To summarize the findings, cumulative maps were generated over all social features to highlight areas where AQ scores significantly modulated neural responses to multiple social features. Finally, the generalizability across fMRI datasets was assessed.

## 2 Methods

### 2.1 Participants

A total of 104 participants were recruited for the fMRI study. The exclusion criteria included a history of neurological or psychiatric disorders, alcohol or substance abuse, BMI under 20 or over 30, current use of psychoactive medications and the standard MRI exclusion criteria. Dataset-specific quality control procedures led to the exclusion of different participants in each dataset. For the **movie clips** dataset, seven participants were excluded (two due to gradient coil malfunction, two due to structural abnormalities, and three due to visible motion artefacts in the preprocessed fMRI data), resulting in a final sample of 97 participants (50 females; mean age = 31 years; range = 20–57). For the **full movie** dataset, eight participants were excluded (two due to gradient coil malfunction, two due to structural abnormalities, and four due to excessive motion), yielding a final sample of 96 participants (47 females; mean age = 31.5 years; range = 20–57). All subjects provided written informed consent and were compensated for their participation. The study was conducted in accordance with the Helsinki Declaration and was approved by the ethics board of the hospital district of Southwest Finland.

### 2.2 Autistic traits of study sample

Autistic traits were measured using AQ questionnaire (Baron-Cohen et al., 2001). The AQ is a 50-item self-report scale designed to assess the degree of autistic traits across five domains: Imagination, Communication, Attention switching, Social skills, and Attention to detail. Participants responded on a 4-point Likert scale ranging from “definitely agree” to “definitely disagree”, with items scored as 1 or 0 (26 reverse-scored items). Individual items were summed to yield total and subscale scores, with higher scores reflecting higher levels of autistic traits. As the study sample did not contain participants with neuropsychiatric disorders, the AQ questionnaire was used to measure the variation of autistic traits within the normal range. The Cronbach’s α coefficient for subscales was from 0.49 to 0.57 (see **Table S1**).

### 2.3 The movie stimulus

Two sets of validated movie stimuli were utilized in the study, both with variable social and emotional contents (Karjalainen et al., 2017; Lahnakoski et al., 2012; Nummenmaa et al., 2023; Santavirta et al., 2023). For the **movie clips** dataset, participants watched a series of 96 English movie clips (19 min 44 s) extracted from Hollywood movies, while a whole-length Finnish feature-film “Käsky” (70 min 11s) was used in the **full movie** dataset (Louhimies, 2008; Santavirta et al., 2024). Participants were instructed to watch movie stimuli as if they were viewing a movie at cinema or at home. Comparisons across the datasets allow testing how well the findings generalize across different naturalistic stimuli (i.e., from short unrelated movie clips to a full movie with continuous narrative).

### 2.4 Annotated social features

The movie stimuli were dynamically annotated for 126 social features with four second temporal resolution to allow parametric modelling of fMRI data. These feature annotations were collected in a separate behavioral experiment, where participants (**movie clips** dataset: n = 6; **full movie** dataset: n = 5) indicated the extent to which each feature was perceived, on a scale from “absent” to “extremely much” that represented continuous values from 0 to 100. To allow reliable modelling of the independent fMRI samples with the average social feature ratings, it was necessary to identify which features are perceived consistently across individuals and are also present in the stimuli. Only features with an intra-class correlation coefficient (ICC) greater than 0.5 were considered to be rated consistently across participants. The ICC2 model was selected since both subjects and movie clips were assumed to represent random samples from their respective populations and videos A feature was considered to be present often enough for modelling if the averaged rating was over five in at least five rating time points. Based on these criteria, 44 social features were identified as reliable predictors for fMRI modelling. Detailed descriptions of the experimental design and stimuli can be found in previous studies using same movie-watching paradigms (Santavirta et al., 2023; Santavirta et al., 2024). The raw average ratings of 44 features in each dataset are presented in Supplementary **Figures S1** and **S2**.

### 2.5 MRI data acquisition and preprocessing

The MRI data were collected using a Phillips Ingenuity TF PET/MR 3T scanner at Turku PET Centre. Functional volumes were acquired with a T2* - weighted echo - planner imaging (EPI) sequence (TR 2600 ms, TE 30 ms, flip angle 75°, FOV 240 mm, 80 × 80 reconstruction matrix, 62.5 kHz bandwidth, 3.0 mm slice thickness, 45 interleaved axial slices acquired in ascending order without gaps). Structural images were obtained with a T1w sequence (1 mm^3^ resolution, TR 9.8 ms, TE 4.6 ms, flip angle 7°, FOV 256 mm, 256 × 256 reconstruction matrix).

The structural and functional imaging data were preprocessed with fMRIPrep (Esteban et al., 2019) (v 1.3.0). First and last two functional volumes were discarded to exclude the time points before and after the stimulus. T1w reference images were corrected for intensity non-uniformity using N4BiasFieldCorrection (Tustison et al., 2010) (v2.1.0) and skull-stripped using antsBrainExtraction.sh (v2.1.0) and OASIS as template. Brain surfaces were reconstructed with recon-all (FreeSurfer v6.0.1) (Dale et al., 1999), and the brain mask estimated previously was refined with a custom variation of the method to reconcile ANTs-derived and FreeSurfer-derived segmentations of the cortical grey matter of Mindboggle (Klein et al., 2017). Spatial normalization to the ICBM 152 Nonlinear Asymmetrical template version 2009c (Fonov et al., 2009) was performed through nonlinear registration with the antsRegistration (ANTs v2.1.0) (Avants et al., 2008), using brain-extracted versions of both T1w volume and template. Brain tissue segmentation of cerebrospinal fluid, white-matter and grey-matter was performed on the brain-extracted T1w image using FAST (Zhang et al., 2001) (FSL v5.0.9).

Functional data were preprocessed as follows: slice-timing correction was done using 3dTshift and motion-corrected using MCFLIRT. The preprocessed BOLD was then co-registered to the T1w image using bbregister for boundary-based registration with six degrees of freedom. All transformations (i.e., motion-correction, coregistration, and spatial normalization) were concatenated and applied in a single step using antsApplyTransforms (ANTs) and Lanczos interpolation. Automatic removal of motion artifacts was performed using ICA-AROMA, followed by spatial smoothing with an isotropic 6-mm Gaussian kernel.

### 2.6 Modelling of the fMRI data

The fMRI data were analyzed in SPM12 (Wellcome Trust Center for Imaging, London, UK, https://www.fil.ion.ucl.ac.uk/spm/). To investigate how autistic traits modulate the neural responses associated with social perception of different features in the general population, we first modelled the neural responses to the 44 dynamically annotated social features in separate first-level models and then added the participant-specific AQ scores as predictors in the second-level models.

Before first-level modelling, standardized rating time series of the social features were convolved with the canonical HRF. Neural responses to each social feature were estimated with separate independent regression models for each feature. Participant-specific first-level contrasts for social features were then subjected to group-level regression analyses. AQ total score and subscale scores were added as predictors in separate second-level models to identify how different autistic traits modulate the neural responses to social features. Statistically significant clusters were identified using one sample t-tests (voxel-level threshold *p* < 0.05 with cluster-level family-wise error rate (FWE) correction, *p* < 0.05). To summarize the results on the regional level, mean beta weights for 56 bilateral anatomical regions of interest (ROI) defined by the AAL2 atlas (Rolls et al., 2015) were extracted for each participant. Then, second-level modelling of the regional data was conducted similar to the full-volume analysis to estimate the modulatory effect of AQ score on the regional level.

### 2.7 Cumulative maps for AQ score effects

To identify the regions where brain responses to different social features were most consistently associated with autistic traits, we calculated cumulative maps for the AQ modulatory effects. This was achieved by first binarizing (1 = significant effect, 0 = no effect) the statistically thresholded beta coefficient maps for each feature, and then calculating the sum of these binarized maps. These cumulative maps were calculated first separately for the two datasets, highlighting the regions where multiple AQ effects were observed.

To investigate the cross-dataset consistency of cumulative maps, we calculated the minimum number of significant feature responses for each brain area across the datasets, which highlights regions where multiple AQ effects were consistently observed in both datasets. Furthermore, on a regional level, we identified the peak AQ effect in each anatomical region by averaging values above the 95^th^ percentile of the accumulation across all voxels within the region and correlated the peak values across datasets to identify anatomical regions with consistent AQ effects. Peak values were used instead of raw regional averages to make anatomical regions of vastly different sizes more comparable with each other.

## 3 Results

### 3.1 Demographic characteristics

In all participants, AQ total score and Imagination subscale score were significantly higher in males than females (**Table S1**). There were no significant correlations between age and AQ scores, except for Attention to detail (*r* = 0.24, *p* = 0.018). Supplementary **Figure S3** shows the correlations between AQ total score and subscale scores.

### 3.2 AQ-related modulation of neural responses to social features in the Movie clips dataset

Next, we examined how consistently autistic traits were associated with responses to social features across the brain (Figure 2). This analysis revealed both general and specific patterns of associations between social perception and AQ dimensions in the **movie clips** dataset, with most consistent effects being observed in the temporal cortex. AQ total score was positively associated with neural responses in the STG and precuneus, while negative association was observed in the fusiform gyrus, orbitofrontal cortex (OFC), insula, MTG, and anterior cingulate cortex (ACC). Regarding the AQ subscales, Attention to detail showed the strongest effects followed by Imagination. Specifically, Attention to detail subscale score showed positive association with neural responses in the frontal pole, ACC, and cuneus gyrus, and negative association was observed in the inferior temporal gyrus (ITG) and lateral occipital cortex (LOC). Imagination subscale score was positively associated with neural responses in the STG, MTG, fusiform gyrus, precuneus, LOC and supramarginal gyrus (SMG), and negative association was observed in the parahippocampal gyrus. Communication subscale score was also positively linked to neural responses in the STG, MTG, and SMG, and negative link was observed in the insula. Additionally, positive relationships between Attention switching score and neural responses were found in the STG, LOC, and posterior cingulate cortex (PCC), while negative relationships were found in the OFC and frontal pole. Unlike the other subscales, Social skills score showed selectively negative association with BOLD responses in subcortical regions (i.e., hippocampus and amygdala), frontal pole, and supplementary motor area (SMA), and positive association was observed in the STG and fusiform gyrus.

**Figure 2.**
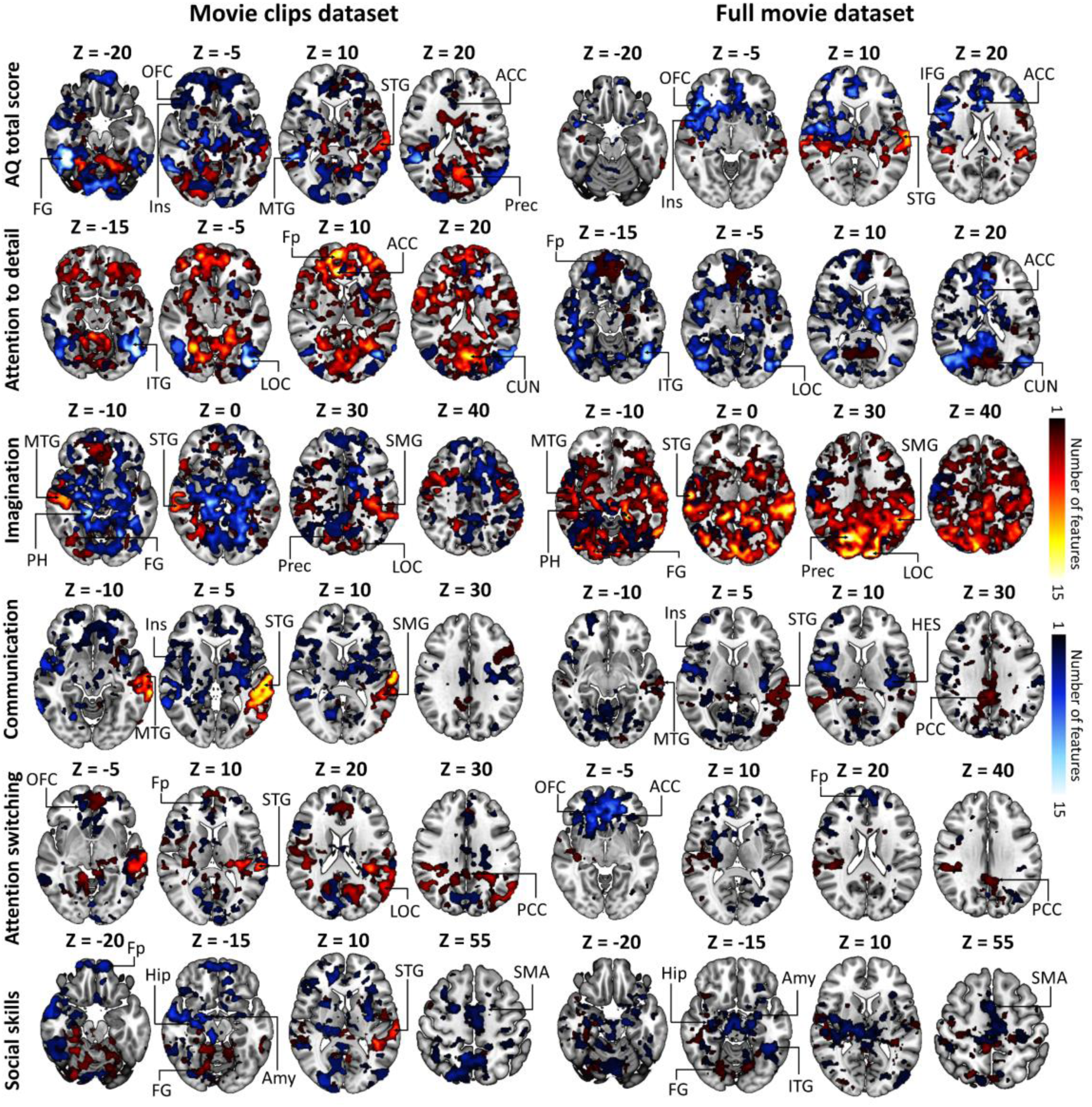
Cumulative maps on AQ-dependent neural responses to multiple social features during movie viewing. Voxel intensities indicate the number of social features (out of 44) where AQ scores were positively (hot colors) or negatively (cold colors) associated with feature-dependent BOLD responses. ACC = anterior cingulate cortex, Amy = amygdala, CUN = cuneus, FG = fusiform gyrus, Fp= frontal pole, Hip = hippocampus, HES = Heschl gyrus, IFG = Inferior frontal gyrus, Ins = Insula, ITG = Inferior temporal gyrus, MTG = middle temporal gyrus, OFC = orbitofrontal cortex, PCC = posterior cingulate cortex, PH = parahippocampal gyrus, PoCG = Postcentral Gyrus, Prec = precuneus, SMA = supplementary motor area, SMG = supramarginal gyrus, STG = superior temporal gyrus.

### 3.3 AQ-related modulation of neural responses to social features in the Full movie dataset

To assess the generalizability of our findings to other movie stimuli, we examined the modulatory effects of AQ in the **full movie** dataset. AQ total score was positively associated with neural responses in the STG, while negative association was observed in the OFC, insula, inferior frontal gyrus (IFG) and ACC. Regarding the AQ subscales, Attention to detail subscale score showed primarily negative association with BOLD responses in the ITG, LOC and ACC, and positive association was observed in the frontal pole and cuneus gyrus. Conversely, Imagination subscale score showed strongly positive association with BOLD response in the fusiform gyrus, STG, MTG, precuneus, LOC and SMG, and negative association was observed in the parahippocampal gyrus. Communication subscale score was negatively associated with neural responses in the insula and Hesch’s gyrus, and positive association was observed in the STG, MTG and PCC. Attention switching score was negatively associated with neural response in the OFC, ACC and frontal pole, while positive association was observed in the PCC. Social skills score showed negative association with BOLD responses in the hippocampus, amygdala, ITG and SMA, and positive association was observed in the fusiform gyrus.

### 3.4 Consistent AQ modulation across datasets

Figure 3 shows a consistent pattern of associations between autistic traits and neural responses to social features across the two datasets. Generally, AQ total score was positively associated with neural responses in the STG, and negative association was observed in the OFC, ACC, and insula. Dimension-specific patterns were also evident: Attention to detail subscale score was negatively linked to neural responses in the LOC, ACC and ITG, and positive association was observed in the cuneus gyrus and frontal pole. Imagination subscale score showed positive association with neural responses in the SMG, STG, MTG, LOC, precuneus and fusiform gyrus, and negative association was observed in the parahippocampal gyrus. Additionally, Communication subscale score was negatively associated with neural responses in insula and positive association was found in the STG and MTG. Attention switching subscale score was negatively associated with neural responses in the OFC and frontal pole, and positive association was observed in the PCC. Social skills subscale score was negatively linked to BOLD responses in the hippocampus, amygdala, and SMA, and positive correlation was observed in the fusiform gyrus.

**Figure 3.**
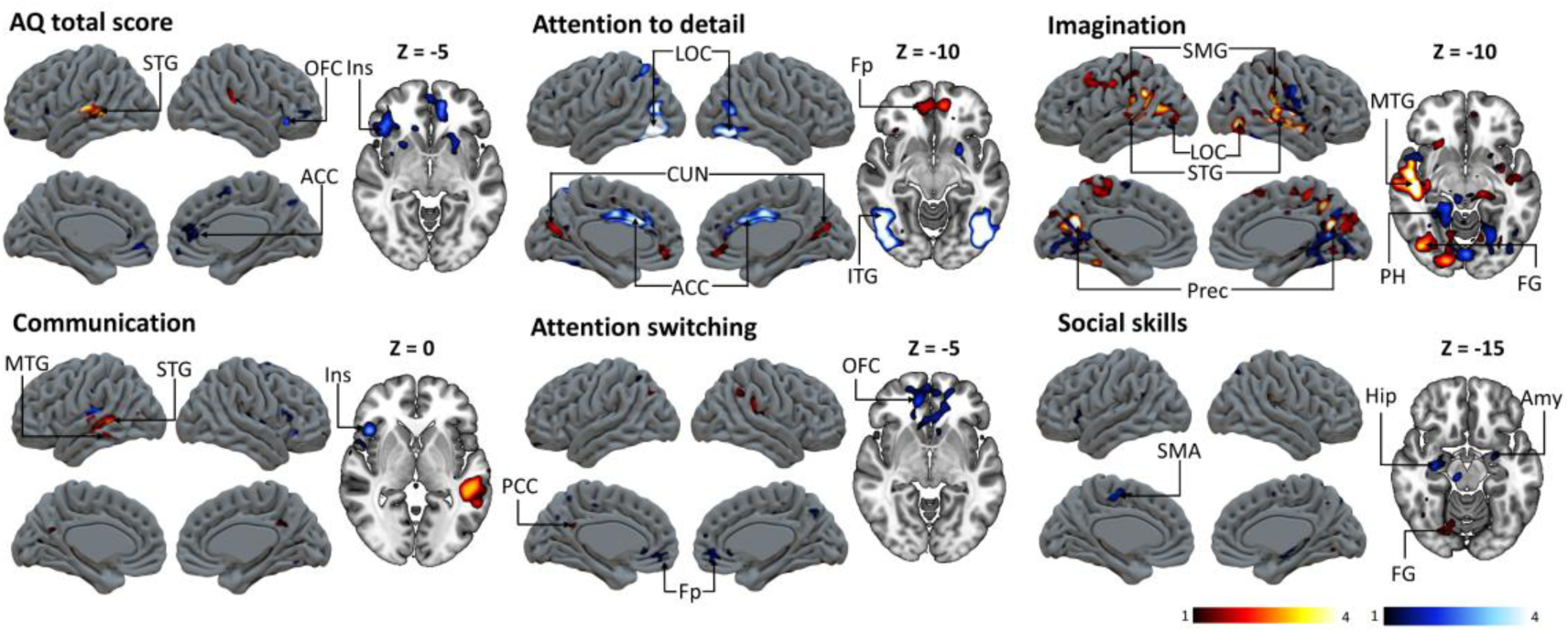
Consistent AQ-related modulation effects across the two datasets. Color bar indicates the range of minimum cumulative value in each brain area where AQ scores showed consistent positive (hot colors) or negative (cold colors) associations with feature-dependent BOLD responses in both datasets. ACC = anterior cingulate cortex, Amy = amygdala, CUN = cuneus, FG = fusiform gyrus, Fp= frontal pole, Hip = hippocampus, Ins = Insula, ITG = Inferior temporal gyrus, MTG = middle temporal gyrus, OFC = orbitofrontal cortex, PH = parahippocampal gyrus, Prec = precuneus, SMA = supplementary motor area, SMG = supramarginal gyrus, STG = superior temporal gyrus.

To assess the spatial similarity of the cumulative responses across two datasets, we conducted correlation analyses with the ROI analysis results. Results revealed differences in consistency across datasets in AQ-related modulation patterns, with Pearson’s *r* values ranging from −0.28 to 0.63 (Figure 4). Positive modulation was most consistent in Attention switching, followed by Communication and Imagination, with overlap in temporal, parietal and frontal cortices. In contrast, for the negative AQ-dependent modulation, cross data consistency was highest for Attention to detail, particularly in occipital, temporal, and frontal cortices. In addition, we assessed the similarity of the broader patterns of AQ dimension and ROI-feature responses across datasets, with the Pearson’s *r* values ranging from 0.08 to 0.22. For the detailed distribution of feature-specific patterns, see Supplementary **Figures S4 and S5**.

**Figure 4.**
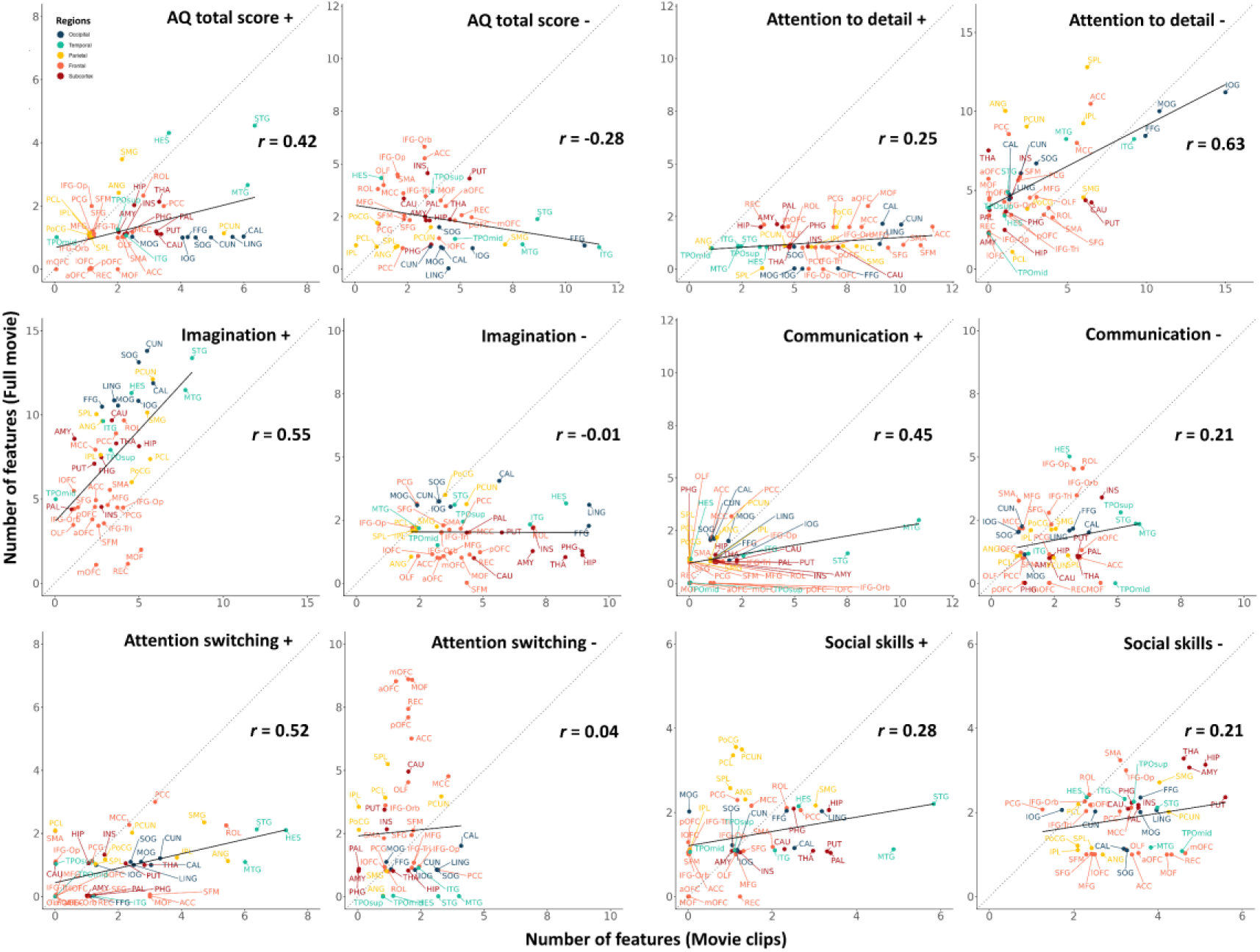
Scatterplot showing the relationship between the Peak AQ effect across the datasets at regional level. The Y-axis (**full movie** dataset) and the X-axis (**movie clips** dataset) represent the number of social features (out of 44) with significant AQ-related modulations. Each point represents an anatomical ROI (see Table S2 for the full name of the regions). For each region, the peak AQ effect was computed as the mean value above the 95^th^ percentile of the accumulation across all voxels within that region. The black regression line depicts the cross-dataset correlation. The gray dashed diagonal marks identical AQ effects across datasets, with the distance of each point from the diagonal reflecting their similarity. The position in relation to the diagonal indicates whether a ROI is more tuned to the movie clips (below diagonal) or full movie (above diagonal).

## 4 Discussion

Our study has three main findings: (1) autistic traits showed replicable general and dimension-specific modulations on the neural responses to a wide range of social perceptual features; (2) autistic traits influenced neural responses to naturalistic audiovisual social stimuli most consistently in the temporal cortex and particularly in the STG across datasets, while AQ subscales showed distinct patterns involving frontal, parietal, occipital, and subcortical regions; and (3) Attention to detail and Imagination subscales were most consistently linked with social perceptual responses, whereas Social skills showed selective pattern involving reduced subcortical activations. These findings indicate that autism-related traits broadly influence neural processing of naturalistic social signals, providing insight into how heterogeneity of autistic symptoms relates to socioemotional processing.

### Brain regions modulated by general autistic traits in social perception

Consistency analysis revealed that the AQ total scores were positively associated with brain responses in the STG, paralleled with negative association in the OFC, ACC and insula (Figure 3).

The STG is associated with speech perception and social processing (Bigler et al., 2007; Lee Masson et al., 2024a; Robins et al., 2009), and is anatomically and functionally connected to the STS (Petrides, 2013; Petrides, 2023), which has been identified as a hub for processing audiovisually presented social and affective information during movie-watching (Lahnakoski et al., 2012; Lee Masson et al., 2024b; Nummenmaa et al., 2012a; Santavirta et al., 2023). These results are hence consistent with prior data highlighting the critical role of temporal cortex in social perception during naturalistic viewing. Moreover, our results align with previous findings in showing that individual differences in autistic traits modulate neural responses particularly in the brain regions involved in social perception (Chen et al., 2023; Nummenmaa et al., 2012a; Puglia and Morris, 2017), broadly similarly as in individuals with ASD (Le Gall and Iakimova, 2018; Peng et al., 2020). Moreover, our data extends these findings to show that autistic traits modulate neural responses to viewing naturalistic social scenes.

The OFC is involved in integrating sensory, emotional, and reward-related information, and in guiding social behavior according to the anticipated value of social cues (Rempel-Clower, 2007; Rushworth et al., 2011). Hypoactivation in OFC is consistently observed in ASD (Patriquin et al., 2016), which may reflect the diminished ability to detect social salience and reduced social motivation.

The ACC is associated with emotion regulation (Etkin et al., 2006) and empathy (Arioli et al., 2021), and dysfunctions in ACC are significantly associated with social cognition impairments in ASD (Di Martino et al., 2009). The insula is associated with emotion recognition and interoception (Terasawa et al., 2021) and its hypoactivation may lead to deficits in emotional processing in ASD (Di Martino et al., 2009). Therefore, AQ-dependent responses to social signals in the STG, OFC, ACC and insula support the notion that the neural mechanisms underlying social perception in ASD extend across a broader spectrum that includes typically developing individuals.

### Brain regions modulated by specific autistic traits in social perception

Several AQ dimension-specific modulations on the neural responses to social perceptual features were replicable across datasets. Scores on the Attention to detail subscale were negatively associated with neural responses in the ACC, ITG and LOC, and positive association was found in the frontal pole and cuneus gyrus (Figure 3). The cuneus gyrus plays a role in basic visual processing, such as encoding position, local orientation, edges and colors (Cohen, 2011; Palejwala et al., 2021), whereas the ITG and LOC are part of the ventral visual stream (Kravitz et al., 2013), responsible for higher-level visual processing, including holistic or global shape, face, and object representations (Cichy et al., 2011; Miyahara et al., 2013), indicating an enhanced sensitivity to local features alongside diminished engagement in global visual integration. On the other hand, the frontal pole is involved in goal-directed persistence (Burgess et al., 2007), and the ACC is well known for its role in conflict monitoring and cognitive control (Agam et al., 2010; Etkin et al., 2006), suggesting that additional neural resources may be recruited to sustain a detail-oriented cognitive style, and reduced flexibility in shifting between different attention strategies.

Imagination subscale score showed the strongest positive association with neural responses in regions located in the parietal and adjacent multimodal association cortices (i.e., STG, MTG, SMG, and precuneus), as well as occipitotemporal regions (i.e., LOC and fusiform gyrus). These regions have been implicated in visual mental imagery (Pearson, 2019; Spagna et al., 2024; Suchan et al., 2002) and also involved in visual perception. In contrast, negative association was observed in the parahippocampal gyrus that is related to autobiographical memory retrieval and scene construction (Aminoff et al., 2013; Mouga et al., 2022). This pattern aligns with a recent fMRI meta-analysis of visual mental imagery, which emphasizes that imagination engages a distributed frontal-temporal-occipital network: prefrontal regions initiate the process by recruiting semantic information from the temporal lobe and activating high-level occipitotemporal areas, while attention and working memory systems sustain and manipulate the reconstructed mental images (Spagna et al., 2021). Within this framework, difficulties to imagine in individuals with autistic traits may result from an imbalance between over-recruitment of semantic and perceptual networks and under-recruitment of memory-based scene construction mechanisms.

The Communication subscale score was significantly correlated with both decreased and increased activation in different brain regions. Lower communicative ability was linked to decreased activation in the insula and increased activation in the STG and MTG. Decreased activation in the insula may reflect impaired integration of visual and auditory signals (Bamiou et al., 2003), social motivation (Clements et al., 2018) and emotion processing (Thom et al., 2014), while increased activation in the STG and MTG may be a compensatory response as an attempt to understand conversions and manage social interactions (Landsiedel and Koldewyn, 2023; Leonard and Chang, 2014). Regarding the Attention switching subscale, we observed a negative association between deficits in attention switching and neural responses in the frontal pole and OFC, both of which are key frontal regions implicated in cognitive flexibility, such as attentional shifting (Hampshire and Owen, 2006). Meanwhile, increased activation associated with attentional inflexibility was observed in the PCC, a core region of the default mode network that is implicated in internally directed attention (Leech et al., 2011). The AQ-dependent modulation in the present study may reflect an impaired capacity to intentionally shift between external perceptual features and internal cognitive control, and to adapt behavior in response to changing environmental demands (Dajani and Uddin, 2015), which is correlated with increased restricted interests and repetitive behaviors in ASD (Faja and Nelson Darling, 2019; Miller et al., 2015).

Finally, social skills, ranging from perception of social cues to more complex competencies like theory of mind (Morrison et al., 2017; Soto-Icaza et al., 2015), are recognized as a foundation for the development of social functioning (Klin et al., 2002). In the present study, Social skills subscale score was observed to be negatively linked to BOLD responses in the limbic systems (i.e., hippocampus and amygdala), and dysfunction in these areas has been observed in ASD during social cognition, including eye contact (Stuart et al., 2023; Tottenham et al., 2014), and emotion recognition (Schultz, 2005; Wei et al., 2025), and inferring other’s mental states (Arioli et al., 2021). The SMA exhibited reduced activation with lower social skills scores, which aligns with its role in observing, imitating, and monitoring movements (Delbruck et al., 2019). Such decreased activation may indicate a reduced ability to accurately perceive nonverbal communicative behaviors and to adaptively adjust behaviors under various social contexts. The fusiform gyrus is a core region for face processing, such as identity recognition and expression recognition (Floris et al., 2025). Although earlier studies indicated that individuals with ASD exhibit hypoactivation in this region when processing emotional face stimuli (Scherf et al., 2015; Wei et al., 2025), our results revealed increased activation in the fusiform gyrus associated with lower social skills scores. This finding aligns with prior evidence showing that the fusiform gyrus is hyper-responsive in individuals with ASD during response inhibition to emotional faces (Duerden et al., 2013; Shafritz et al., 2015). One possible explanation is that individuals with diminished social skills may recruit additional neural resources to detect and interpret socially salient facial cues during movie watching.

Altogether, our study revealed neural alterations associated with autism-like traits in the general population, which resemble patterns observed in ASD, and thus support the notion of behavioral heterogeneity in autism (Dufour et al., 2013; Pina-Camacho et al., 2012; Weston, 2019). Moreover, our findings align with previous work that demonstrated increased idiosyncratic responses in brain areas important for social cognition and their correlations with autistic traits in ASD during movie viewing (Hasson et al., 2009; Lyons et al., 2020; Salmi et al., 2013; Turner et al., 2025), suggesting that individual differences in autism-like traits among the general populations modulate the way people perceive complex and naturalistic social interactions.

## 5 Limitations

As our study only involved healthy volunteers, and the AQ scores were distributed within the normal range, we cannot tell whether the similar patterns of AQ-dependent compensatory modulations to naturalistic social features extend to individuals clinically diagnosed with ASD. Future studies could compare the presently observed findings with those stemming from individuals with clinically diagnosed ASD. Second, we did not measure whether participants had socially relevant difficulties in perceiving or inferring the stimulus scenes during the experiments, which would have given more insights into the possible reasons why the neural responses in certain regions were modulated by AQ scores. Third, GLM and correlation analyses showed discrepancies in AQ-related modulations in neural responses to social features across datasets (Figures 2 and **4**), which may partly reflect variability in the naturalistic stimulus episodes used. Nonetheless, the convergent results from the consistency analysis identified regions most reliably engaged in perceiving social features across diverse naturalistic contexts. Finally, autism frequently co-occurs with many other conditions, like alexithymia and anxiety symptoms. Future studies should disentangle the unique and overlapping contributions of these conditions to the brain basis of socioemotional processing.

## 6 Conclusions

In sum, we demonstrated that individual differences in autism-like traits in neurotypical individuals are associated with neural responses to social features in regions involved in multisensory, social, and cognitive functions during naturalistic movie viewing. Most consistent effects across features and datasets were observed in the temporal cortices. These associations indicate that neural alterations underlying social perception in ASD extend across a broader spectrum that includes typically developing individuals. Because these neural alterations varied across distinct dimensions of autism-related traits, they provide neurobiological support for the heterogeneous nature of ASD.

## Acknowledgements

This work was supported by Signe of Ane Gyllenberg’s Stiftelse, Jane and Aatos Erkko foundation, Research Council of Finland (grant #350416), and China Scholarship Council (202408330143).

## Supplementary materials

**Figure S1.**
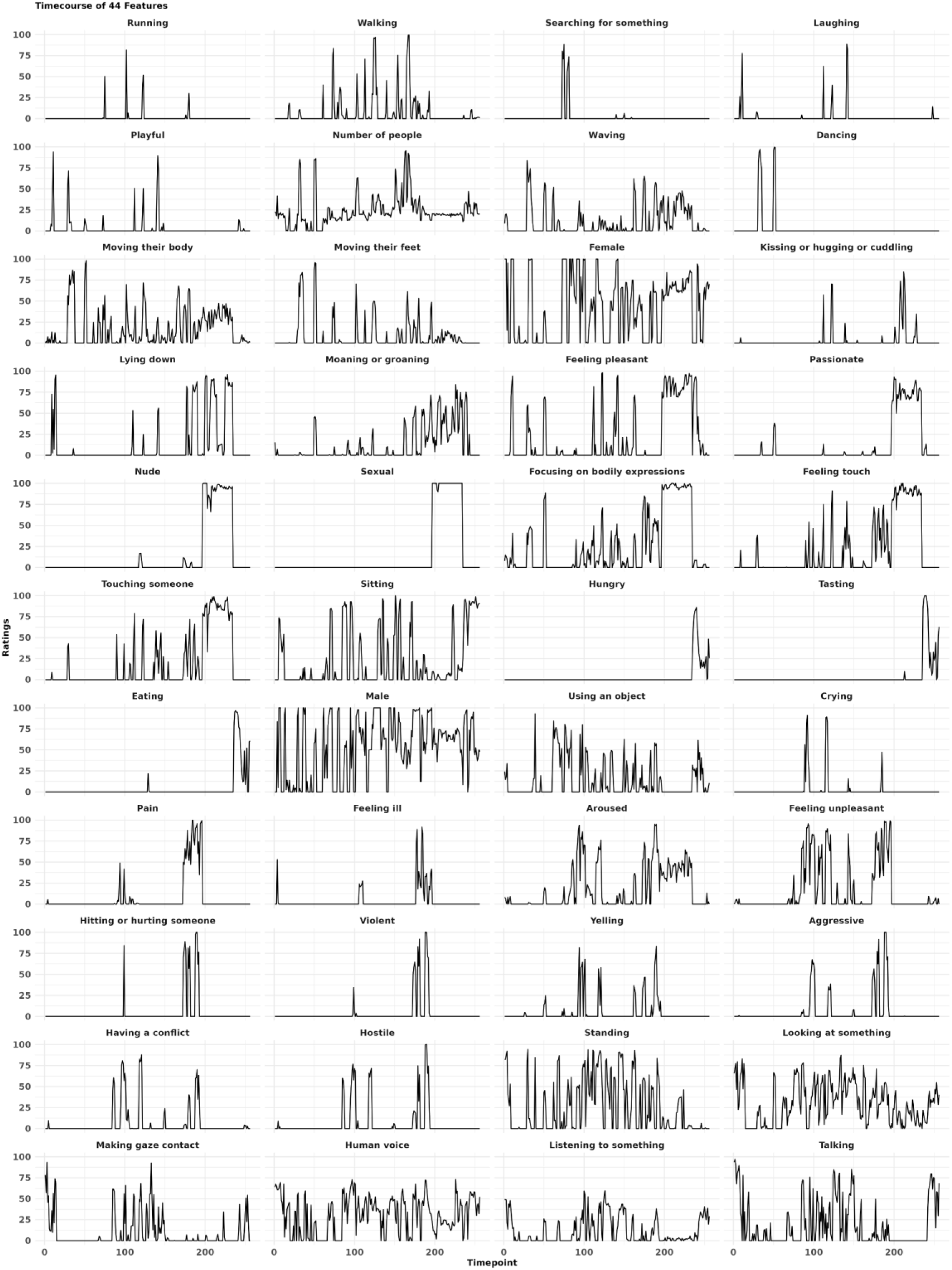
Raw averaged ratings for 44 social features in the **movie clips** dataset.

**Figure S2.**
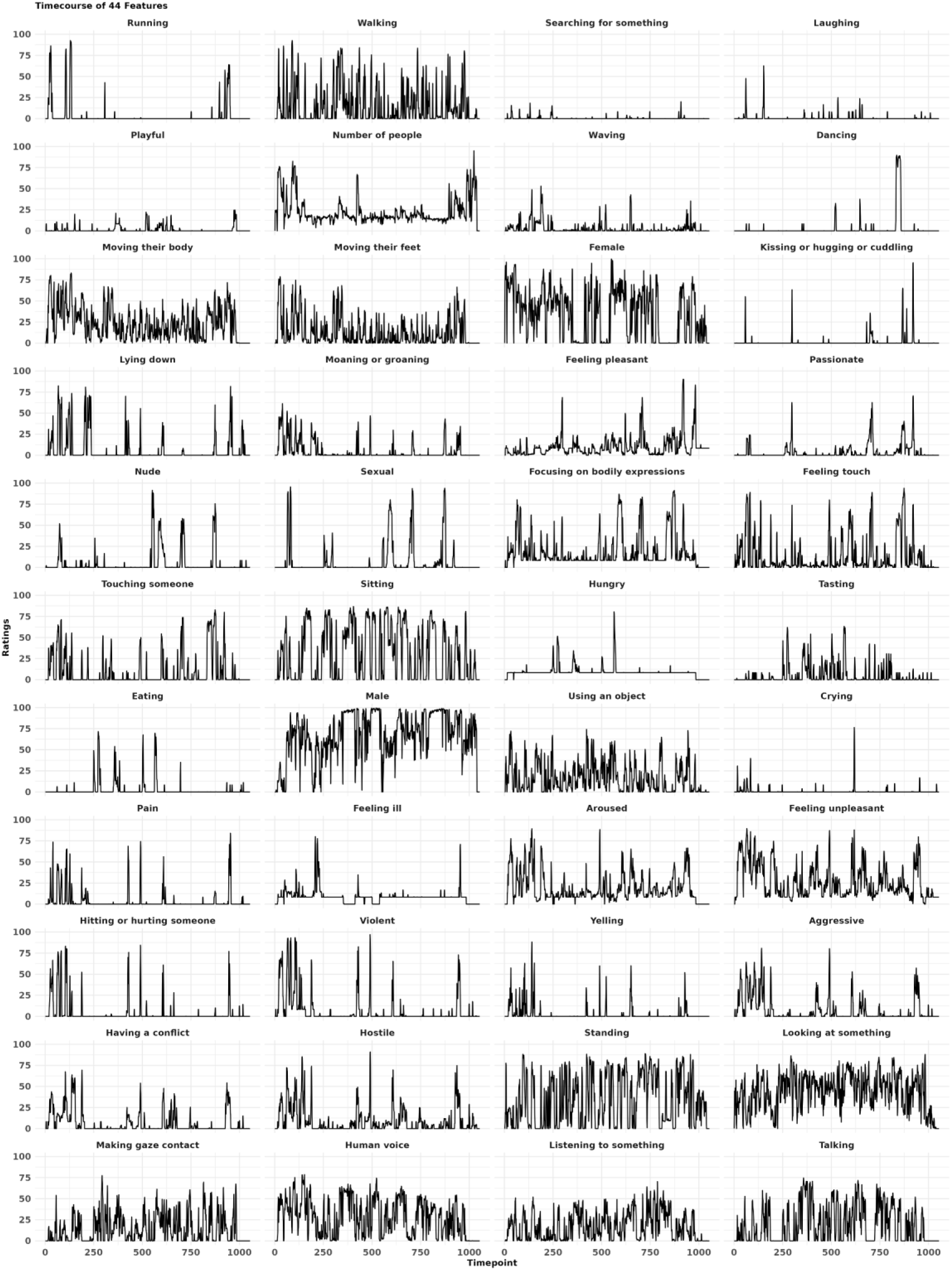
Raw averaged ratings for 44 social features in the full movie dataset.

**Figure S3.**
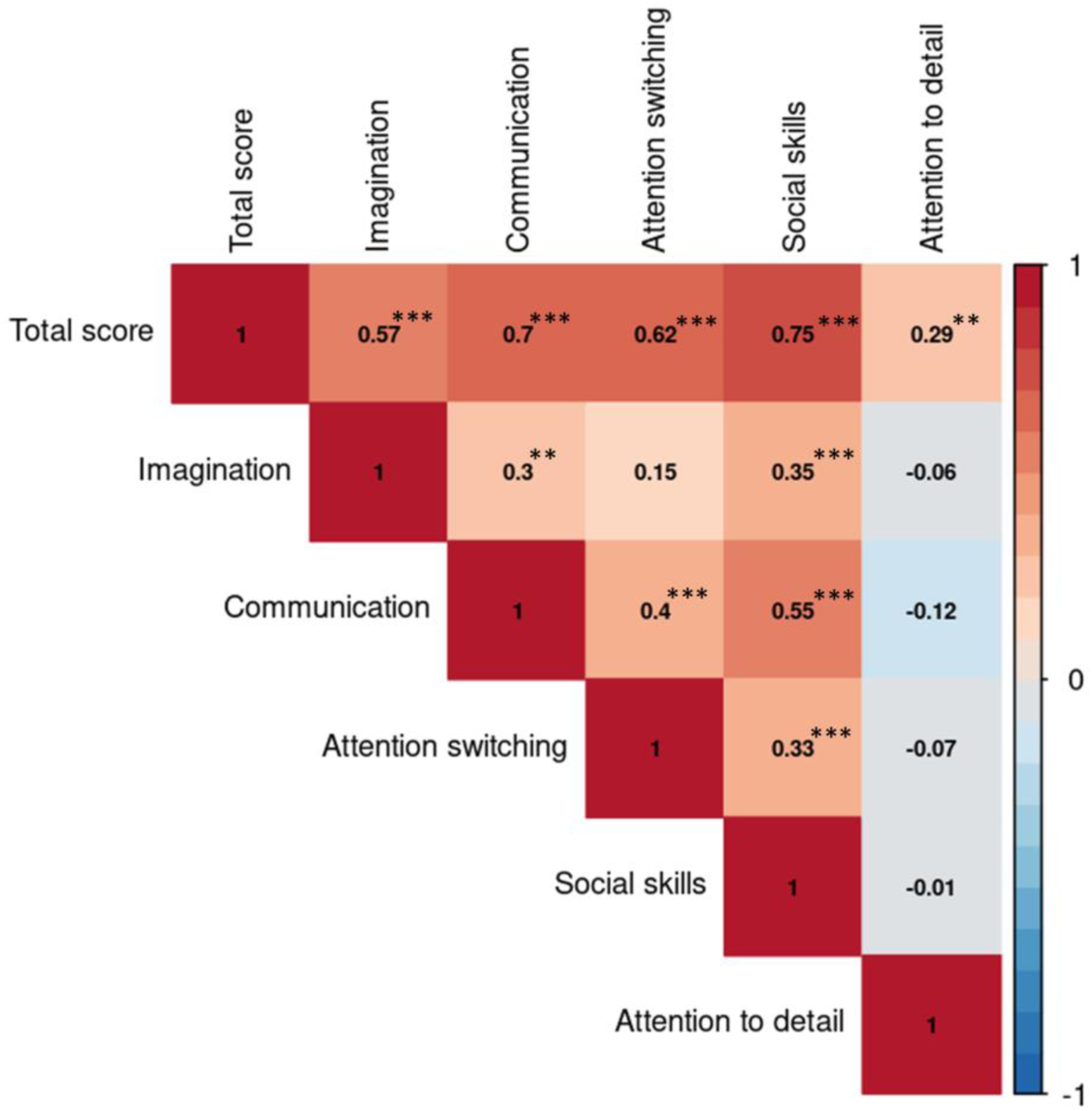
Correlations between AQ total and subscale scores in all participants (N = 100). ** indicates *p* < 0.01; *** indicates *p* < 0.001.

**Figure S4.**
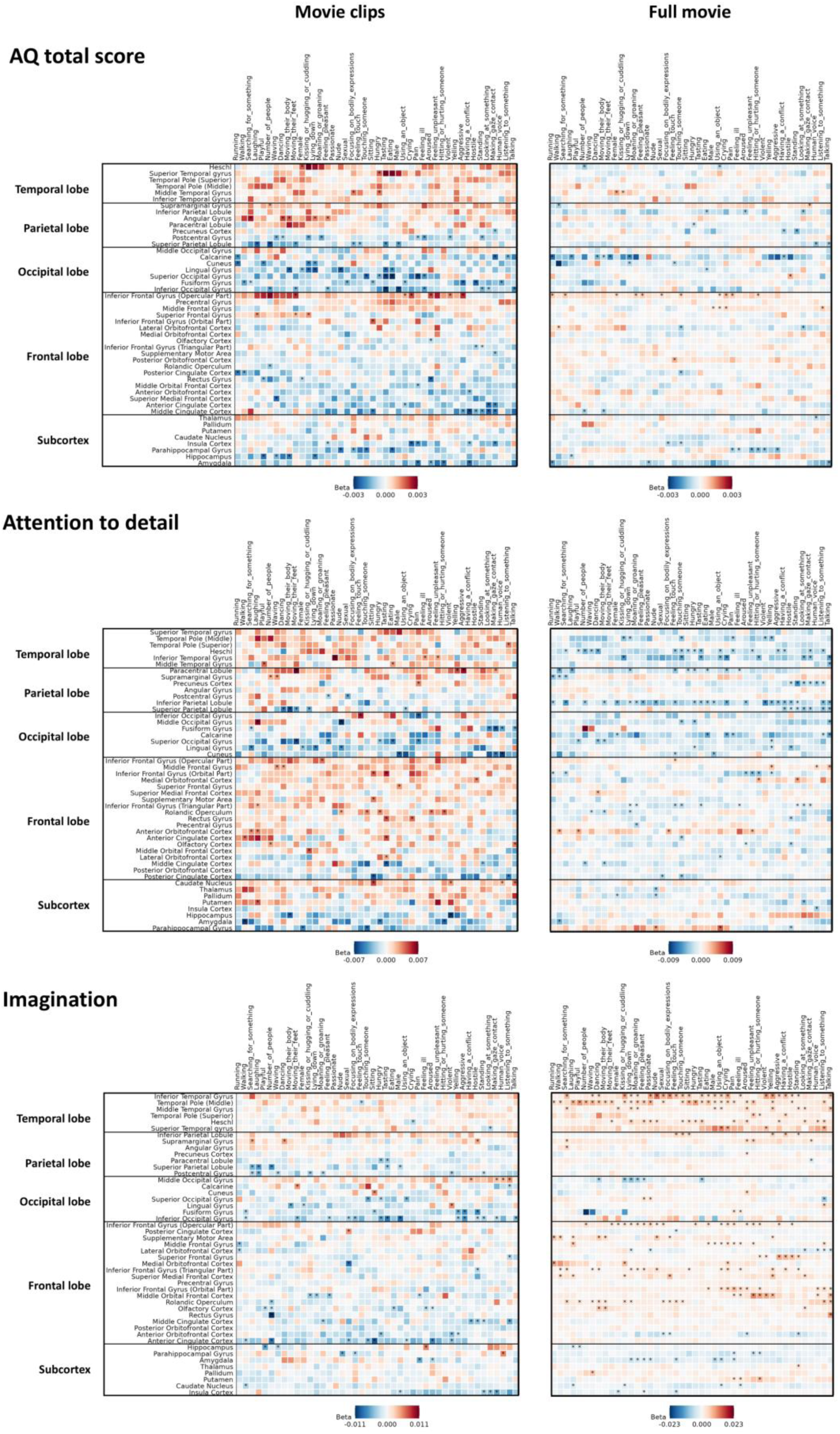
Regression patterns between AQ dimensions (i.e., AQ total score, Attention to detail, and Imagination) and ROI-feature responses. Asterisks indicate statistical significance (*p* < 0.05).

**Figure S5.**
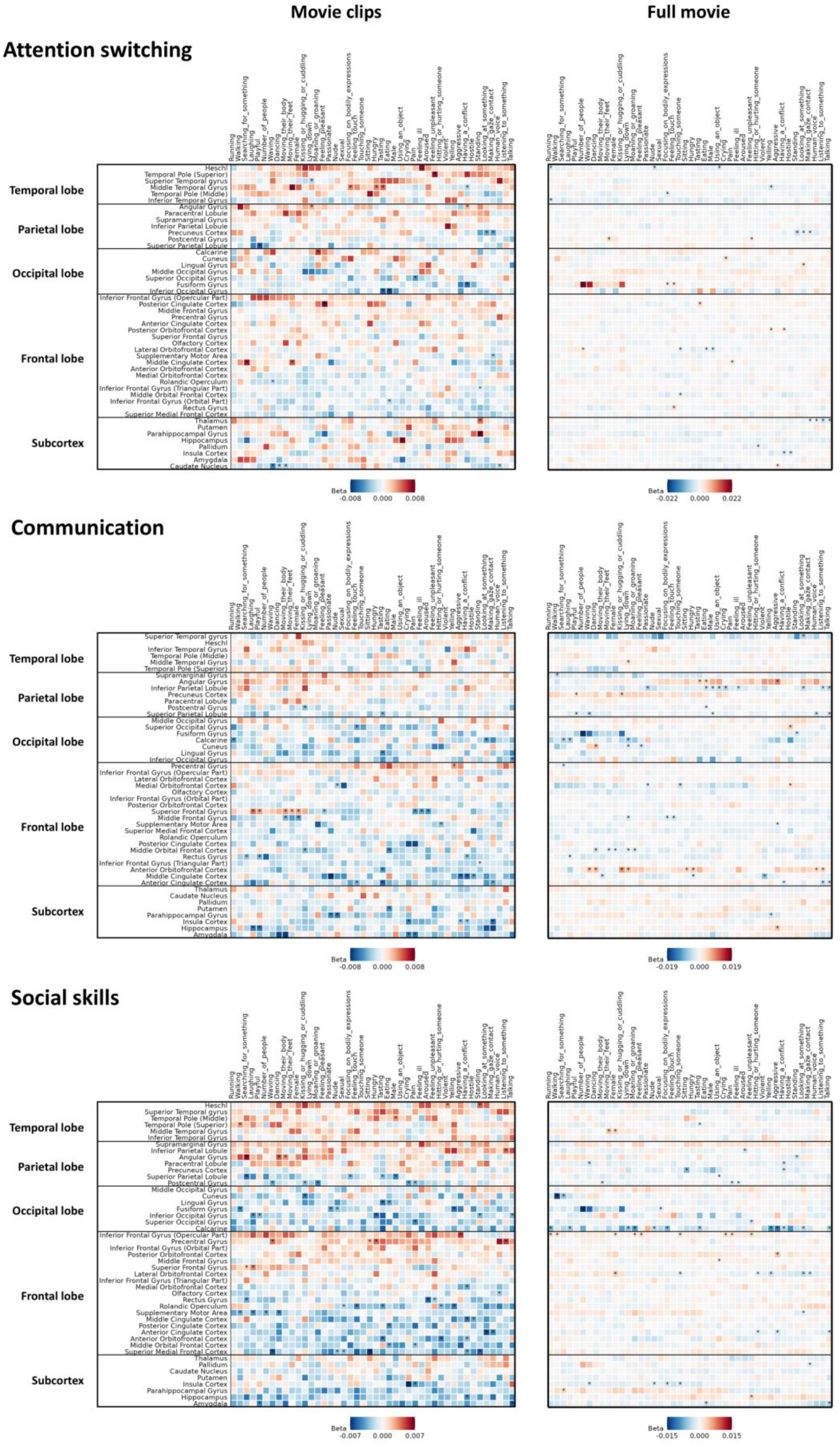
Regression patterns between remaining AQ dimensions (i.e., Attention switching, Communication, and Social skills) and ROI-feature responses. Asterisks indicate statistical significance (*p* < 0.05)

**Table S1.**
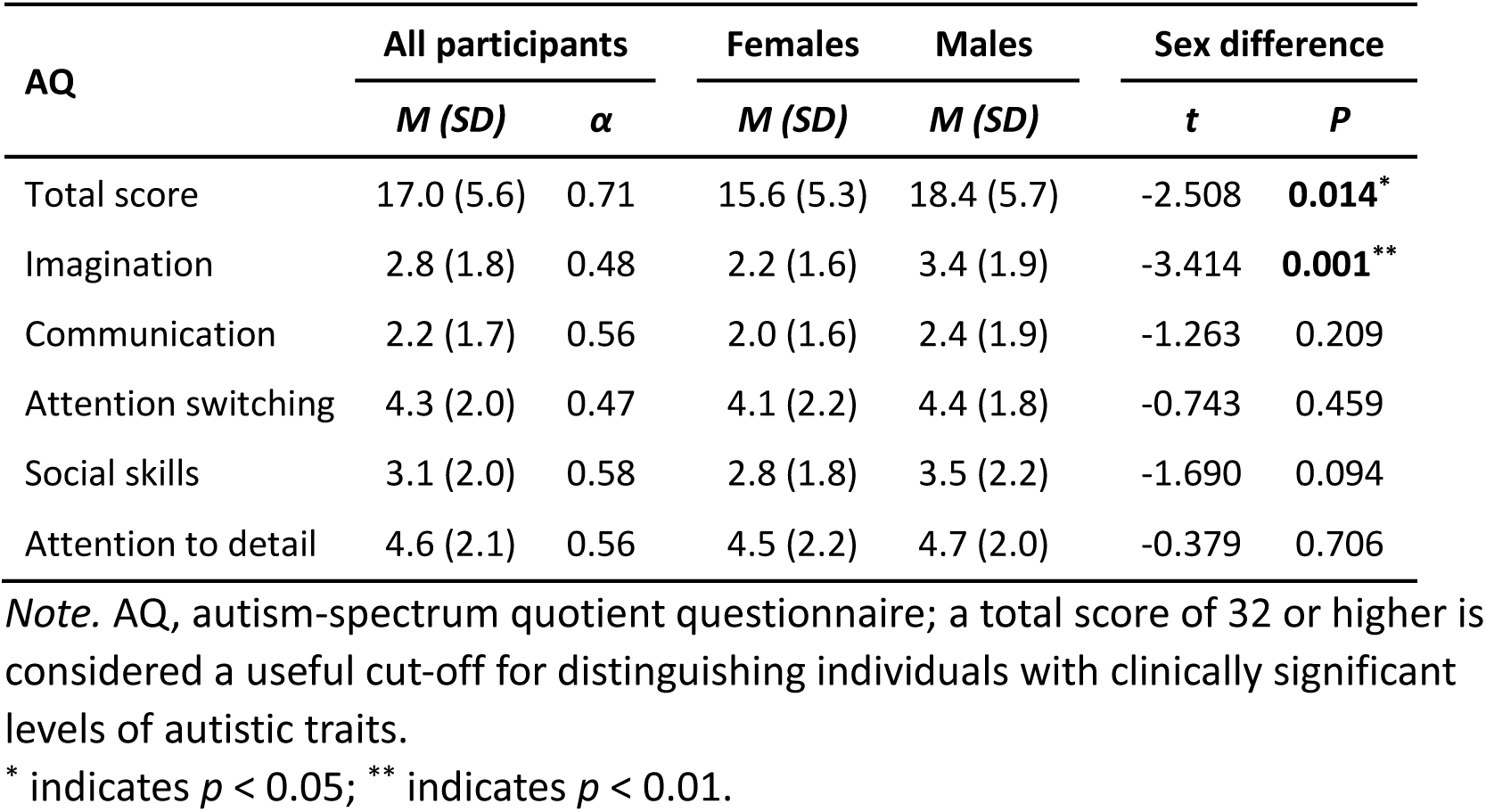
Scores and the Cronbach’s alpha reliability coefficients of the AQ (all participants, N = 100).

**Table S2.** Brain regions presented in Figure 4.

| Abbreviation | Full Name |
| --- | --- |
| PCG | Precentral Gyrus |
| SFG | Superior Frontal Gyrus |
| MFG | Middle Frontal Gyrus |
| IFG-Op | Inferior Frontal Gyrus (Opercular Part) |
| IFG-Tri | Inferior Frontal Gyrus (Triangular Part) |
| IFG-Orb | Inferior Frontal Gyrus (Orbital Part) |
| ROL | Rolandic Operculum |
| SMA | Supplementary Motor Area |
| OLF | Olfactory Cortex |
| SFM | Superior Medial Frontal Cortex |
| MOF | Middle Orbital Frontal Cortex |
| REC | Rectus Gyrus |
| mOFC | Medial Orbitofrontal Cortex |
| aOFC | Anterior Orbitofrontal Cortex |
| pOFC | Posterior Orbitofrontal Cortex |
| IOFC | Lateral Orbitofrontal Cortex |
| INS | Insular Cortex |
| ACC | Anterior Cingulate Cortex |
| MCC | Middle Cingulate Cortex |
| PCC | Posterior Cingulate Cortex |
| HIP | Hippocampus |
| PHG | Parahippocampal Gyrus |
| AMY | Amygdala |
| CAL | Calcarine |
| CUN | Cuneus |
| LING | Lingual Gyrus |
| SOG | Superior Occipital Gyrus |
| MOG | Middle Occipital Gyrus |
| IOG | Inferior Occipital Gyrus |
| FFG | Fusiform Gyrus |
| PoCG | Postcentral Gyrus |
| SPL | Superior Parietal Lobule |
| IPL | Inferior Parietal Lobule |
| SMG | Supramarginal Gyrus |
| ANG | Angular Gyrus |
| PCUN | Precuneus |
| PCL | Paracentral Lobule |
| CAU | Caudate Nucleus |
| PUT | Putamen |
| PAL | Pallidum |
| THA | Thalamus |
| HES | Heschl |
| STG | Superior Temporal Gyrus |
| TPOsup | Temporal Pole (Superior) |
| MTG | Middle Temporal Gyrus |
| TPOmid | Temporal Pole (Middle) |
| ITG | Inferior Temporal Gyrus |

## Notes

### Competing Interest Statement

The authors have declared no competing interest.

### Summary of Updates

Minor corrections: removed an extra comma from the keyword list and corrected the title of Table S2. No other changes were made.

## Reference

Agam, Y., Joseph, R.M., Barton, J.J., Manoach, D.S., 2010. Reduced cognitive control of response inhibition by the anterior cingulate cortex in autism spectrum disorders. Neuroimage 52, 336–347. 10.1016/j.neuroimage.2010.04.010

Alkhaldi, R.S., Sheppard, E., Ellerby, Z., Burdett, E.R.R., Mitchell, P., 2024. The relationship between autistic traits, expressiveness, readability and social perceptions. PloS One 19, e0301003.

Allison, T., Puce, A., McCarthy, G., 2000. Social perception from visual cues: role of the STS region. Trends in Cognitive Sciences 4, 267–278. 10.1016/s1364-6613(00)01501-1

Aminoff, E.M., Kveraga, K., Bar, M., 2013. The role of the parahippocampal cortex in cognition. Trends in Cognitive Sciences 17, 379–390. 10.1016/j.tics.2013.06.009

Amodio, D.M., Frith, C.D., 2006. Meeting of minds: the medial frontal cortex and social cognition. Nature Reviews: Neuroscience 7, 268–277. 10.1038/nrn1884

Arioli, M., Cattaneo, Z., Ricciardi, E., Canessa, N., 2021. Overlapping and specific neural correlates for empathizing, affective mentalizing, and cognitive mentalizing: A coordinate-based meta-analytic study. Human Brain Mapping 42, 4777–4804. 10.1002/hbm.25570

Association, A.P., 2013. Diagnostic and statistical manual of mental disorders. American psychiatric association.

Avants, B.B., Epstein, C.L., Grossman, M., Gee, J.C., 2008. Symmetric diffeomorphic image registration with cross-correlation: evaluating automated labeling of elderly and neurodegenerative brain. Medical Image Analysis 12, 26–41. 10.1016/j.media.2007.06.004

Bamiou, D.E., Musiek, F.E., Luxon, L.M., 2003. The insula (Island of Reil) and its role in auditory processing. Literature review. Brain Research: Brain Research Reviews 42, 143–154. 10.1016/s0165-0173(03)00172-3

Baron-Cohen, S., Wheelwright, S., Skinner, R., Martin, J., Clubley, E., 2001. The Autism-Spectrum Quotient (AQ): Evidence from Asperger syndrome/high-functioning autism, males and females, scientists and mathematicians. Journal of Autism and Developmental Disorders 31, 5–17. 10.1023/a:1005653411471

Bigler, E.D., Mortensen, S., Neeley, E.S., Ozonoff, S., Krasny, L., Johnson, M., Lu, J., Provencal, S.L., McMahon, W., Lainhart, J.E., 2007. Superior temporal gyrus, language function, and autism. Developmental Neuropsychology 31, 217–238. 10.1080/87565640701190841

Burgess, P.W., Dumontheil, I., Gilbert, S.J., 2007. The gateway hypothesis of rostral prefrontal cortex (area 10) function. Trends in Cognitive Sciences 11, 290–298. 10.1016/j.tics.2007.05.004

Chen, T., Gau, S.S., Wu, Y.Y., Chou, T.L., 2023. Neural substrates of theory of mind in adults with autism spectrum disorder: An fMRI study of the social animation task. Journal of the Formosan Medical Association 122, 621–628. 10.1016/j.jfma.2022.10.009

Chita-Tegmark, M., 2016. Social attention in ASD: A review and meta-analysis of eye-tracking studies. Research in Developmental Disabilities 48, 79–93. 10.1016/j.ridd.2015.10.011

Cichy, R.M., Chen, Y., Haynes, J.D., 2011. Encoding the identity and location of objects in human LOC. Neuroimage 54, 2297–2307. 10.1016/j.neuroimage.2010.09.044

Clements, C.C., Zoltowski, A.R., Yankowitz, L.D., Yerys, B.E., Schultz, R.T., Herrington, J.D., 2018. Evaluation of the Social Motivation Hypothesis of Autism: A Systematic Review and Meta-analysis. JAMA Psychiatry 75, 797–808. 10.1001/jamapsychiatry.2018.1100

Cohen, R.A., 2011. Cuneus. Encyclopedia of clinical neuropsychology. Springer, pp. 756–757.

Constantino, J.N., Davis, S.A., Todd, R.D., Schindler, M.K., Gross, M.M., Brophy, S.L., Metzger, L.M., Shoushtari, C.S., Splinter, R., Reich, W., 2003. Validation of a brief quantitative measure of autistic traits: comparison of the social responsiveness scale with the autism diagnostic interview-revised. Journal of Autism and Developmental Disorders 33, 427–433. 10.1023/a:1025014929212

Corluka, N., Laycock, R., 2024. The influence of dynamism and expression intensity on face emotion recognition in individuals with autistic traits. Cognition & Emotion 38, 635–644. 10.1080/02699931.2024.2314982

Dajani, D.R., Uddin, L.Q., 2015. Demystifying cognitive flexibility: Implications for clinical and developmental neuroscience. Trends in Neurosciences 38, 571–578. 10.1016/j.tins.2015.07.003

Dale, A.M., Fischl, B., Sereno, M.I., 1999. Cortical surface-based analysis. I. Segmentation and surface reconstruction. Neuroimage 9, 179–194. 10.1006/nimg.1998.0395

Deen, B., Koldewyn, K., Kanwisher, N., Saxe, R., 2015. Functional Organization of Social Perception and Cognition in the Superior Temporal Sulcus. Cerebral Cortex 25, 4596–4609. 10.1093/cercor/bhv111

Di Martino, A., Ross, K., Uddin, L.Q., Sklar, A.B., Castellanos, F.X., Milham, M.P., 2009. Functional brain correlates of social and nonsocial processes in autism spectrum disorders: an activation likelihood estimation meta-analysis. Biological Psychiatry 65, 63–74. 10.1016/j.biopsych.2008.09.022

Duerden, E.G., Taylor, M.J., Soorya, L.V., Wang, T., Fan, J., Anagnostou, E., 2013. Neural correlates of inhibition of socially relevant stimuli in adults with autism spectrum disorder. Brain Research 1533, 80–90. 10.1016/j.brainres.2013.08.021

Dufour, N., Redcay, E., Young, L., Mavros, P.L., Moran, J.M., Triantafyllou, C., Gabrieli, J.D.E., Saxe, R., 2013. Similar Brain Activation during False Belief Tasks in a Large Sample of Adults with and without Autism. PloS One 8. 10.1371/journal.pone.0075468

Esteban, O., Markiewicz, C.J., Blair, R.W., Moodie, C.A., Isik, A.I., Erramuzpe, A., Kent, J.D., Goncalves, M., DuPre, E., Snyder, M., Oya, H., Ghosh, S.S., Wright, J., Durnez, J., Poldrack, R.A., Gorgolewski, K.J., 2019. fMRIPrep: a robust preprocessing pipeline for functional MRI. Nature Methods 16, 111–116. 10.1038/s41592-018-0235-4

Etkin, A., Egner, T., Peraza, D.M., Kandel, E.R., Hirsch, J., 2006. Resolving emotional conflict: a role for the rostral anterior cingulate cortex in modulating activity in the amygdala. Neuron 51, 871–882. 10.1016/j.neuron.2006.07.029

Faja, S., Nelson Darling, L., 2019. Variation in restricted and repetitive behaviors and interests relates to inhibitory control and shifting in children with autism spectrum disorder. Autism 23, 1262–1272. 10.1177/1362361318804192

Floris, D.L., Llera, A., Zabihi, M., Moessnang, C., Jones, E.J.H., Mason, L., Haartsen, R., Holz, N.E., Mei, T., Elleaume, C., Vieira, B.H., Pretzsch, C.M., Forde, N.J., Baumeister, S., Dell’Acqua, F., Durston, S., Banaschewski, T., Ecker, C., Holt, R.J., Baron-Cohen, S., Bourgeron, T., Charman, T., Loth, E., Murphy, D.G.M., Buitelaar, J.K., Beckmann, C.F., Holz, N.E., Forde, N.J., Banaschewski, T., Langer, N., 2025. A multimodal neural signature of face processing in autism within the fusiform gyrus. Nature Mental Health 3. 10.1038/s44220-024-00349-4

Fonov, V.S., Evans, A.C., McKinstry, R.C., Almli, C.R., Collins, D., 2009. Unbiased nonlinear average age-appropriate brain templates from birth to adulthood. Neuroimage 47, S102.

Hampshire, A., Owen, A.M., 2006. Fractionating attentional control using event-related fMRI. Cerebral Cortex 16, 1679–1689. 10.1093/cercor/bhj116

Hasson, U., Avidan, G., Gelbard, H., Vallines, I., Harel, M., Minshew, N., Behrmann, M., 2009. Shared and idiosyncratic cortical activation patterns in autism revealed under continuous real-life viewing conditions. Autism Research 2, 220–231. 10.1002/aur.89

Hasson, U., Nir, Y., Levy, I., Fuhrmann, G., Malach, R., 2004. Intersubject synchronization of cortical activity during natural vision. Science 303, 1634–1640. 10.1126/science.1089506

Hudson, M., Santavirta, S., Putkinen, V., Seppala, K., Sun, L., Karjalainen, T., Karlsson, H.K., Hirvonen, J., Nummenmaa, L., 2023. Neural responses to biological motion distinguish autistic and schizotypal traits. Social Cognitive and Affective Neuroscience 18. 10.1093/scan/nsad011

Huth, A.G., Nishimoto, S., Vu, A.T., Gallant, J.L., 2012. A continuous semantic space describes the representation of thousands of object and action categories across the human brain. Neuron 76, 1210–1224. 10.1016/j.neuron.2012.10.014

Karjalainen, T., Karlsson, H.K., Lahnakoski, J.M., Glerean, E., Nuutila, P., Jaaskelainen, I.P., Hari, R., Sams, M., Nummenmaa, L., 2017. Dissociable Roles of Cerebral mu-Opioid and Type 2 Dopamine Receptors in Vicarious Pain: A Combined PET-fMRI Study. Cerebral Cortex 27, 4257–4266. 10.1093/cercor/bhx129

Klein, A., Ghosh, S.S., Bao, F.S., Giard, J., Hame, Y., Stavsky, E., Lee, N., Rossa, B., Reuter, M., Chaibub Neto, E., Keshavan, A., 2017. Mindboggling morphometry of human brains. PLoS Computational Biology 13, e1005350. 10.1371/journal.pcbi.1005350

Klin, A., Jones, W., Schultz, R., Volkmar, F., Cohen, D., 2002. Visual fixation patterns during viewing of naturalistic social situations as predictors of social competence in individuals with autism. Archives of General Psychiatry 59, 809–816. 10.1001/archpsyc.59.9.809

Kravitz, D.J., Saleem, K.S., Baker, C.I., Ungerleider, L.G., Mishkin, M., 2013. The ventral visual pathway: an expanded neural framework for the processing of object quality. Trends in Cognitive Sciences 17, 26–49. 10.1016/j.tics.2012.10.011

Lahnakoski, J.M., Glerean, E., Salmi, J., Jääskeläinen, I., Sams, M., Hari, R., Nummenmaa, L., 2012. Naturalistic fMRI mapping reveals superior temporal sulcus as the hub for the distributed brain network for social perception. Frontiers in Human Neuroscience 6. 10.3389/fnhum.2012.00233

Landsiedel, J., Koldewyn, K., 2023. Auditory dyadic interactions through the “eye” of the social brain: How visual is the posterior STS interaction region? Imaging Neuroscience 1. 10.1162/imag_a_00003

Le, B.l., Huang, Y.h., Wang, L.l., Hu, H.x., Wang, X., Wang, Y., Wang, Y., Huang, J., Lui, S.S., Chan, R.C., 2024. Individuals with high levels of autistic traits exhibit impaired cognitive but not affective theory of mind and empathy. PsyCh Journal 13, 486–493.

Le Gall, E., Iakimova, G., 2018. [Social cognition in schizophrenia and autism spectrum disorder: Points of convergence and functional differences]. Encephale 44, 523–537. 10.1016/j.encep.2018.03.004

Lee Masson, H., Chang, L., Isik, L., 2024a. Multidimensional neural representations of social features during movie viewing. Social Cognitive and Affective Neuroscience 19. 10.1093/scan/nsae030

Lee Masson, H., Chen, J., Isik, L., 2024b. A shared neural code for perceiving and remembering social interactions in the human superior temporal sulcus. Neuropsychologia 196, 108823. 10.1016/j.neuropsychologia.2024.108823

Leech, R., Kamourieh, S., Beckmann, C.F., Sharp, D.J., 2011. Fractionating the default mode network: distinct contributions of the ventral and dorsal posterior cingulate cortex to cognitive control. Journal of Neuroscience 31, 3217–3224. 10.1523/JNEUROSCI.5626-10.2011

Leonard, M.K., Chang, E.F., 2014. Dynamic speech representations in the human temporal lobe. Trends in Cognitive Sciences 18, 472–479. 10.1016/j.tics.2014.05.001

Limanowski, J., Blankenburg, F., 2016. That’s not quite me: limb ownership encoding in the brain. Social Cognitive and Affective Neuroscience 11, 1130–1140.

Louhimies, A., 2008. Käsky.

Lyons, K.M., Stevenson, R.A., Owen, A.M., Stojanoski, B., 2020. Examining the relationship between measures of autistic traits and neural synchrony during movies in children with and without autism. Neuroimage-Clinical 28. 10.1016/j.nicl.2020.102477

Miller, H.L., Ragozzino, M.E., Cook, E.H., Sweeney, J.A., Mosconi, M.W., 2015. Cognitive Set Shifting Deficits and Their Relationship to Repetitive Behaviors in Autism Spectrum Disorder. Journal of Autism and Developmental Disorders 45, 805–815. 10.1007/s10803-014-2244-1

Miyahara, M., Harada, T., Ruffman, T., Sadato, N., Iidaka, T., 2013. Functional connectivity between amygdala and facial regions involved in recognition of facial threat. Social Cognitive and Affective Neuroscience 8, 181–189. 10.1093/scan/nsr085

Morrison, K.E., Pinkham, A.E., Penn, D.L., Kelsven, S., Ludwig, K., Sasson, N.J., 2017. Distinct profiles of social skill in adults with autism spectrum disorder and schizophrenia. Autism Research 10, 878–887. 10.1002/aur.1734

Mouga, S., Duarte, I.C., Cafe, C., Sousa, D., Duque, F., Oliveira, G., Castelo-Branco, M., 2022. Parahippocampal deactivation and hyperactivation of central executive, saliency and social cognition networks in autism spectrum disorder. Journal of Neurodevelopmental Disorders 14, 9. 10.1186/s11689-022-09417-1

Nummenmaa, L., Engell, A.D., von dem Hagen, E., Henson, R.N., Calder, A.J., 2012a. Autism spectrum traits predict the neural response to eye gaze in typical individuals. Neuroimage 59, 3356–3363. 10.1016/j.neuroimage.2011.10.075

Nummenmaa, L., Glerean, E., Viinikainen, M., Jaaskelainen, I.P., Hari, R., Sams, M., 2012b. Emotions promote social interaction by synchronizing brain activity across individuals. Proceedings of the National Academy of Sciences of the United States of America 109, 9599–9604. 10.1073/pnas.1206095109

Nummenmaa, L., Malen, T., Nazari-Farsani, S., Seppala, K., Sun, L., Santavirta, S., Karlsson, H.K., Hudson, M., Hirvonen, J., Sams, M., Scott, S., Putkinen, V., 2023. Decoding brain basis of laughter and crying in natural scenes. Neuroimage 273, 120082. 10.1016/j.neuroimage.2023.120082

Palejwala, A.H., Dadario, N.B., Young, I.M., O’Connor, K., Briggs, R.G., Conner, A.K., O’Donoghue, D.L., Sughrue, M.E., 2021. Anatomy and White Matter Connections of the Lingual Gyrus and Cuneus. World Neurosurgery 151, e426–e437. 10.1016/j.wneu.2021.04.050

Pantelis, P.C., Byrge, L., Tyszka, J.M., Adolphs, R., Kennedy, D.P., 2015. A specific hypoactivation of right temporo-parietal junction/posterior superior temporal sulcus in response to socially awkward situations in autism. Social Cognitive and Affective Neuroscience 10, 1348–1356. 10.1093/scan/nsv021

Patriquin, M.A., DeRamus, T., Libero, L.E., Laird, A., Kana, R.K., 2016. Neuroanatomical and neurofunctional markers of social cognition in autism spectrum disorder. Human Brain Mapping 37, 3957–3978. 10.1002/hbm.23288

Pearson, J., 2019. The human imagination: the cognitive neuroscience of visual mental imagery. Nature Reviews: Neuroscience 20, 624–634. 10.1038/s41583-019-0202-9

Pelphrey, K.A., Morris, J.P., McCarthy, G., 2005. Neural basis of eye gaze processing deficits in autism. Brain 128, 1038–1048. 10.1093/brain/awh404

Peng, Z.W., Chen, J.R., Jin, L.L., Han, H.Y., Dong, C.J., Guo, Y., Kong, X.J., Wan, G.B., Wei, Z., 2020. Social brain dysfunctionality in individuals with autism spectrum disorder and their first-degree relatives: An activation likelihood estimation meta-analysis. Psychiatry Research-Neuroimaging 298. 10.1016/j.pscychresns.2020.111063

Petrides, M., 2013. Neuroanatomy of language regions of the human brain. Academic Press.

Petrides, M., 2023. On the evolution of polysensory superior temporal sulcus and middle temporal gyrus: A key component of the semantic system in the human brain. Journal of Comparative Neurology 531, 1987–1995. 10.1002/cne.25521

Pina-Camacho, L., Villero, S., Fraguas, D., Boada, L., Janssen, J., Navas-Sanchez, F.J., Mayoral, M., Llorente, C., Arango, C., Parellada, M., 2012. Autism spectrum disorder: does neuroimaging support the DSM-5 proposal for a symptom dyad? A systematic review of functional magnetic resonance imaging and diffusion tensor imaging studies. Journal of Autism and Developmental Disorders 42, 1326–1341.

Puglia, M.H., Morris, J.P., 2017. Neural Response to Biological Motion in Healthy Adults Varies as a Function of Autistic-Like Traits. Frontiers in Neuroscience 11. 10.3389/fnins.2017.00404

Redcay, E., 2008. The superior temporal sulcus performs a common function for social and speech perception: implications for the emergence of autism. Neurosci Biobehav Rev 32, 123–142. 10.1016/j.neubiorev.2007.06.004

Rempel-Clower, N.L., 2007. Role of orbitofrontal cortex connections in emotion. Annals of the New York Academy of Sciences 1121, 72–86. 10.1196/annals.1401.026

Riddiford, J.A., Enticott, P.G., Lavale, A., Gurvich, C., 2022. Gaze and social functioning associations in autism spectrum disorder: A systematic review and meta-analysis. Autism Research 15, 1380–1446.

Robins, D.L., Hunyadi, E., Schultz, R.T., 2009. Superior temporal activation in response to dynamic audio-visual emotional cues. Brain and Cognition 69, 269–278. 10.1016/j.bandc.2008.08.007

Rolls, E.T., Joliot, M., Tzourio-Mazoyer, N., 2015. Implementation of a new parcellation of the orbitofrontal cortex in the automated anatomical labeling atlas. Neuroimage 122, 1–5. 10.1016/j.neuroimage.2015.07.075

Ropar, D., Greenfield, K., Smith, A.D., Carey, M., Newport, R., 2018. Body representation difficulties in children and adolescents with autism may be due to delayed development of visuo-tactile temporal binding. Developmental Cognitive Neuroscience 29, 78–85. 10.1016/j.dcn.2017.04.007

Rushworth, M.F., Noonan, M.P., Boorman, E.D., Walton, M.E., Behrens, T.E., 2011. Frontal cortex and reward-guided learning and decision-making. Neuron 70, 1054–1069. 10.1016/j.neuron.2011.05.014

Salmi, J., Roine, U., Glerean, E., Lahnakoski, J., Nieminen-von Wendt, T., Tani, P., Leppämäki, S., Nummenmaa, L., Jääskeläinen, I.P., Carlson, S., 2013. The brains of high functioning autistic individuals do not synchronize with those of others. NeuroImage: Clinical 3, 489–497.

Santavirta, S., Karjalainen, T., Nazari-Farsani, S., Hudson, M., Putkinen, V., Seppala, K., Sun, L., Glerean, E., Hirvonen, J., Karlsson, H.K., Nummenmaa, L., 2023. Functional organization of social perception networks in the human brain. Neuroimage 272, 120025. 10.1016/j.neuroimage.2023.120025

Santavirta, S., Malen, T., Erdemli, A., Nummenmaa, L., 2024. A taxonomy for human social perception: Data-driven modeling with cinematic stimuli. Journal of Personality and Social Psychology 127, 1146–1171. 10.1037/pspa0000415

Scherf, K.S., Elbich, D., Minshew, N., Behrmann, M., 2015. Individual differences in symptom severity and behavior predict neural activation during face processing in adolescents with autism. Neuroimage Clin 7, 53–67. 10.1016/j.nicl.2014.11.003

Schultz, R.T., 2005. Developmental deficits in social perception in autism: the role of the amygdala and fusiform face area. International Journal of Developmental Neuroscience 23, 125–141. 10.1016/j.ijdevneu.2004.12.012

Shafritz, K.M., Bregman, J.D., Ikuta, T., Szeszko, P.R., 2015. Neural systems mediating decision-making and response inhibition for social and nonsocial stimuli in autism. Progress in Neuro-Psychopharmacology and Biological Psychiatry 60, 112–120. 10.1016/j.pnpbp.2015.03.001

Skjegstad, C.L., Trevor, C., Swanborough, H., Roswandowitz, C., Mokros, A., Habermeyer, E., Fruhholz, S., 2022. Psychopathic and autistic traits differentially influence the neural mechanisms of social cognition from communication signals. Transl Psychiatry 12, 494. 10.1038/s41398-022-02260-x

Soto-Icaza, P., Aboitiz, F., Billeke, P., 2015. Development of social skills in children: neural and behavioral evidence for the elaboration of cognitive models. Frontiers in Neuroscience 9, 333. 10.3389/fnins.2015.00333

Spagna, A., Hajhajate, D., Liu, J., Bartolomeo, P., 2021. Visual mental imagery engages the left fusiform gyrus, but not the early visual cortex: A meta-analysis of neuroimaging evidence. Neurosci Biobehav Rev 122, 201–217. 10.1016/j.neubiorev.2020.12.029

Spagna, A., Heidenry, Z., Miselevich, M., Lambert, C., Eisenstadt, B.E., Tremblay, L., Liu, Z., Liu, J., Bartolomeo, P., 2024. Visual mental imagery: Evidence for a heterarchical neural architecture. Phys Life Rev 48, 113–131. 10.1016/j.plrev.2023.12.012

Stuart, N., Whitehouse, A., Palermo, R., Bothe, E., Badcock, N., 2023. Eye Gaze in Autism Spectrum Disorder: A Review of Neural Evidence for the Eye Avoidance Hypothesis. Journal of Autism and Developmental Disorders 53, 1884–1905. 10.1007/s10803-022-05443-z

Suchan, B., Yaguez, L., Wunderlich, G., Canavan, A.G., Herzog, H., Tellmann, L., Homberg, V., Seitz, R.J., 2002. Neural correlates of visuospatial imagery. Behavioural Brain Research 131, 163–168. 10.1016/s0166-4328(01)00373-4

Terasawa, Y., Motomura, K., Natsume, A., Iijima, K., Chalise, L., Sugiura, J., Yamamoto, H., Koyama, K., Wakabayashi, T., Umeda, S., 2021. Effects of insular resection on interactions between cardiac interoception and emotion recognition. Cortex 137, 271–281. 10.1016/j.cortex.2021.01.011

Thom, N.J., Johnson, D.C., Flagan, T., Simmons, A.N., Kotturi, S.A., Van Orden, K.F., Potterat, E.G., Swain, J.L., Paulus, M.P., 2014. Detecting emotion in others: increased insula and decreased medial prefrontal cortex activation during emotion processing in elite adventure racers. Social Cognitive and Affective Neuroscience 9, 225–231. 10.1093/scan/nss127

Tottenham, N., Hertzig, M.E., Gillespie-Lynch, K., Gilhooly, T., Millner, A.J., Casey, B.J., 2014. Elevated amygdala response to faces and gaze aversion in autism spectrum disorder. Social Cognitive and Affective Neuroscience 9, 106–117. 10.1093/scan/nst050

Turner, J.M., Byrge, L., Richardson, H., Galdi, P., Kennedy, D.P., Kliemann, D., 2025. Social inference brain networks in autistic adults during movie-viewing: functional specialization and heterogeneity. Molecular Autism 16, 42. 10.1186/s13229-025-00669-x

Tustison, N.J., Avants, B.B., Cook, P.A., Zheng, Y., Egan, A., Yushkevich, P.A., Gee, J.C., 2010. N4ITK: improved N3 bias correction. IEEE Transactions on Medical Imaging 29, 1310–1320. 10.1109/TMI.2010.2046908

van den Heuvel, M.P., Sporns, O., 2011. Rich-club organization of the human connectome. Journal of Neuroscience 31, 15775–15786. 10.1523/JNEUROSCI.3539-11.2011

Voos, A.C., Pelphrey, K.A., Kaiser, M.D., 2013. Autistic traits are associated with diminished neural response to affective touch. Social Cognitive and Affective Neuroscience 8, 378–386. 10.1093/scan/nss009

Wei, L., Zhou, M., Hu, P., Jia, S., Zhong, S., 2025. Abnormal brain activation in autism spectrum disorder during negative emotion processing: A meta-analysis of functional neuroimaging studies. Journal of Psychiatric Research 185, 1–10. 10.1016/j.jpsychires.2025.03.032

Weston, C.S.E., 2019. Four Social Brain Regions, Their Dysfunctions, and Sequelae, Extensively Explain Autism Spectrum Disorder Symptomatology. Brain Sciences 9. 10.3390/brainsci9060130

Zhang, Y.Y., Brady, M., Smith, S., 2001. Segmentation of brain MR images through a hidden Markov random field model and the expectation-maximization algorithm. IEEE Transactions on Medical Imaging 20, 45–57. 10.1109/42.906424

